# Region-specific microRNA alterations in marmosets carrying SLC6A4 polymorphisms are associated with anxiety-like behavior

**DOI:** 10.1101/2021.04.08.437842

**Authors:** Natalia Popa, Dipankar Bachar, Angela C. Roberts, Andrea M. Santangelo, Eduardo Gascon

**Author notes:** Co-senior authors (Andrea M. Santangelo, Eduardo Gascon). Corresponding author: Eduardo Gascon, Institut de Neurosciences de la Timone, Aix-Marseille Université; CNRS UMR7289; 27, Boulevard Jean Moulin, 13005 Marseille (France).

## Abstract

Psychiatric diseases such as depression and anxiety are multifactorial conditions, highly prevalent in western societies. Human studies have identified a number of high-risk genetic variants for these diseases. Among them, polymorphisms in the promoter region of the serotonin transporter gene (SLC6A4) have attracted much attention. However, due to the paucity of experimental models, molecular alterations induced by these genetic variants and how they correlate to behavioral deficits have not been examined. Marmosets have emerged as a powerful model in translational neuroscience to investigate molecular underpinnings of complex behaviors. Here, we took advantage of naturally occurring genetic polymorphisms in marmoset *SLC6A4* gene that have been linked to anxiety-like behaviors. Using FACS-sorted cells from different brain regions, we revealed that marmosets bearing different *SLC6A4* variants exhibit distinct microRNAs signatures in a region of the prefrontal cortex whose activity has been consistently altered in patients with depression/anxiety. We also identified DCC, a gene previously linked to these diseases, as a downstream target of the dysregulated microRNAs. Significantly, we showed that levels of both microRNAs and DCC in this region were highly correlated to anxiety-like behaviors as well as to the response to citalopram, a selective serotonin re-uptake inhibitor and widely prescribed anti-depressant. Our findings establish links between genetic variants, molecular modifications in specific cortical regions and complex behavioral/pharmacological responses, providing new insights into gene-behavior relationships underlying human psychopathology.

## INTRODUCTION

Stress-related disorders such as depression and anxiety are common, highly debilitating and burdensome conditions whose incidence has dramatically increased during the COVID-19 pandemic [1]. Despite multiple therapeutic options, a large proportion of patients show no clinical improvement after treatment. Indeed, only about one third of depressed patients respond to serotonin reuptake inhibitors (SSRIs), the most widely prescribed antidepressants (ADs) [2].

The pathophysiology of psychiatric diseases is complex and genetic factors are thought to play a major role [3]. Indeed, human association studies have pinpointed the links between a number of genetic variants and an increased risk of developing such diseases. Among them, polymorphisms in the SLC6A4, the gene encoding the serotonin transporter, are of particular interest for several reasons. First, since their initial discovery in the 90s [4], multiple studies have linked these polymorphisms with depression/anxiety [5]. Second, later work supported the interaction between stress, a risk factor for developing psychiatric conditions, and SLC6A4 polymorphisms in depression [6, 7]. Third, serotonin neurotransmission is particularly prominent in the regions of the ventro-medial prefrontal cortex (vmPFC) that are dysregulated in patients [8, 9]. Finally, these polymorphisms have also been associated with poor treatment response to ADs [10].

The SLC6A4 gene is located in human chromosome 17q11-12. Although variants have been found across the gene, most studies have focused on the so-called serotonin-transporter linked promoter region (5-HTTLPR) [11, 12]. This region is around 1,5 kb upstream of the first exon and contains a variable number of repeats. Importantly, this particular arrangement of SLC6A4 promoter is conserved in primates but not in rodents (Suppl. Fig. 1). A short allele containing 14 repeats (*5-HTT*LPR S) and a long allele (*5-HTT*LPR L) comprising 16 repeats have been widely documented. Most studies have reported that *5*-HTTLPR S results in lower mRNA and protein levels of the serotonin transporter and, as a consequence, a reduction of circulating serotonin (from the synapse to the presynaptic terminal) [12, 13] It has been proposed that this reduction in serotonin has important downstream functional effects in several brain areas including vmPFC [14]. It has also been linked to a higher risk of psychopathology, modulation of stressful events in depression [4, 6, 15] and impaired response to ADs [10].

In support of human findings, several groups have found an association between psychiatric-like phenotypes and the equivalent short allele in macaques [16]. SLC6A4 variants [17] have also been characterized in the common marmoset, *Callithrix jacchus*, which has emerged as a reference non-human primate model in modern neuroscience [18]. Rather than a difference in repeat number, however, marmoset polymorphisms were discrete sequence substitutions in the promoter region [17]. Importantly, Santangelo et al. found two predominant variants, AC/C/G and CT/T/C, in captivity as well as in the wild. Similar to humans, one of those variants (AC/C/G) resulted in reduced mRNA expression [17]. Marmosets carrying two alleles of this variant exhibited an anxious-like behavior when confronted with a human intruder standing in front of their cage, a poor response to citalopram and altered serotonin receptor 2a binding and mRNA levels in emotion-related brain regions [17, 19]. Together these observations support the notion that SLC6A4 promoter structure as well as its genetic variants are functionally relevant for psychiatric conditions across multiple primate species.

Although 5-HTTLPR polymorphisms influence serotonin transporter levels [4, 11, 13], the mechanisms by which these genetic variants increase the risk of psychopathology are currently unknown. In this regard, microRNAs (miRNAs), a class of short (20-25 nt) non-coding RNAs acting as posttranscriptional repressors of gene expression, are attractive candidates. On one hand, their regulatory potential is vast as most protein coding genes are computationally predicted to be miRNAs targets [20]. On the other hand, previous work has shown selectivity in the relationship between miRNAs and stress-related disorders as well as therapeutic responses [21–23]. Finally, the investigation of miRNAs in psychiatric disorders has gained momentum as accumulating evidence indicates that miRNAs could potentially be used as biomarkers [24].

In order to provide insight into the mechanisms of gene-behavior interactions in primates, we determine whether the distinct *SLC6A4* haplotypes related to trait-like anxiety in marmosets are associated with region-specific alterations in miRNA within the vmPFC. Not only do we reveal such region-specific miRNA alterations related to the *SLC6A4* variants but we also identify DCC, a gene previously implicated in affective disorders, as a downstream target of the deregulated miRNA networks. Such changes are highly correlated with individual behavioral anxiety-like responses. They underscore the intimate link between genetic variants, molecular differences among vmPFC areas and complex behavioral outcomes, of major relevance for our understanding of human psychopathology.

## METHODS

### Subjects and genotyping

Marmosets were bred onsite at the Innes Marmoset Colony (Behavioral and Clinical Neuroscience Institute) and housed as male-female pairs (males were vasectomized). Temperature (24 °C) and humidity (55%) conditions were controlled, and a dawn/dusk-like 12-h period was maintained. They were provided with a balanced diet and water ad libitum. This research has been regulated under the Animals (Scientific Procedures) Act 1986 Amendment Regulations 2012 following ethical review by the University of Cambridge Animal Welfare and Ethical Review Body.

For this study, 6 adult male common marmosets, Callithrix jacchus, (26 ± 2 mo, 413 ± 17 g) balanced for SLC6A4 genotype (Supplementary table 1) were used. All animals had human intruder (HI) test, and snake testing (procedures described previously by [25]) before entering a pharmacological study, which consisted of repeated HI test with acute intramuscular (i.m.) doses of citalopram (behavioral data reported elsewhere [17]) and, after 2 months, with the 5HT2A antagonist M100907 [19].

Genotyping was carried out following the protocol described in [17]. Primers used for sequencing can be found in Supplementary Table 3

### Behavioral testing and quantification

All the behavioral data in this study were collected and reported previously [17, 19, 26]. All the behavioral procedures (see supplementary materials) have been extensively described in the same references as well as in the Supplementary Materials section. Analyzed behaviors are summarized in Supplementary Table 2.

### Pharmacological Manipulation on the intruder test

Animals were injected i.m. with citalopram (2.5 mg/kg), with the selective 5-HT2A antagonist M100907 (Sigma–Aldrich) (0.3 mg/kg) or vehicle (0.01 M PBS-HCl) 25 min before the intruder phase. Procedures for the human intruder test were exactly the same as described above. To avoid habituation to the human intruder across sessions the intruder wore different realistic rubber human masks each session. The experimental design was a latin square randomized by genotype, and masks. Treatment order was the same for all individuals (lower dose, higher dose, and vehicle) with 2 weeks between each session.

### Sample preparation

At the end of the study, animals were premedicated with ketamine hydrochloride before being euthanized with pentobarbital sodium (Dolethal; 1 mL of a 200-mg/mL solution; Merial Animal Health). Brain were dissected, frozen using liquid nitrogen, and then sliced in a cryostat at −20 °C to 200-μm-thick sections. Tissue samples for each target region were excised using punches of 1.0 and 1.5-mm radio length. Eight punches per target region were used in this study (4 from the right hemisphere and 4 from the left hemisphere).

### Nuclei isolation and sorting

8 punches/area/animal were used. Nuclei extraction protocol was adapted from [27]. All steps were performed at 4 °C or on ice. Tissues were homogenized in nuclei isolation buffer (0.32 M Sucrose, 10 mM HEPES pH 8.0, 5 mM CaCl_2_, 3 mM Mg(CH_3_COO)_2_, 0.1 mM EDTA, 1 mM DTT, 0.1% Triton X-100) with a 2 ml Dounce homogenizer by 10 gentle strokes with each pestle and filtered through a 40 μm strainer. After centrifugation, nuclei pellets were resuspended in 1 ml PBS-RI (PBS, 50 U/mL Rnase-OUT Recombinant Ribonuclease Inhibitor (Invitrogen), 1 mM DTT) and fixed by the addition of 3 ml PBS 1.33% paraformaldehyde (Electron Microscopy Sciences) for 30 minutes on ice. Fixed nuclei were spun down, washed with 1 ml PBS 0.1% triton-X-100, pelleted again and resuspended at 10^6^ nuclei per ml in stain/wash buffer (PBS-RI, 0.5% BSA, 0.1% Triton-X-100) containing 2 μg/ml anti-NeuN-alexa-488 antibody (Millipore, MAB377X) and 1 μg/ml Hoechst 33342 (Molecular Probes). After 30 minutes incubation on ice protected from light, nuclei were washed with 2 mL stain/wash buffer and spun down. Finally, stained nuclei were resuspended in 1 ml PBS-RI 0.5% BSA and filtered again through a 40 μm strainer. Nuclei suspensions were maintained on ice protected from light until sorting. Sorting of nuclei was achieved with a MoFlo Astrios EQ Cell sorter (Beckman Coulter). After positive selection of intact Hoechst-positive nuclei and doublets exclusion, all NeuN-positive and NeuN-negative nuclei were separately isolated. Sorted nuclei were collected in refrigerated 2 ml microtubes containing 0.5 ml PBS-RI 0.5% BSA. Finally, nuclei were spun down, supernatants eliminated and pellets were conserved at −80°C untill RNA extraction.

### RNA extraction and reverse transcription

Total RNAs (small and large RNAs) were extracted in one fraction with miRNeasy FFPE kit (Qiagen) following manufacturer’s protocol with minor changes. Briefly, nuclei pellets were lysed in 150 μL PKD buffer and 10 μl proteinase K for 15 minutes at 56°C, then immediately incubated at 80°C for 15 minutes in order to reverse formaldehyde modification of nucleic acids and then immediately incubated 3 minutes on ice. After centrifugation, supernatants were transferred in new 2 ml microtubes and remaining DNA was degraded during a 30 minutes incubation with DNase Booster Buffer and DNase I. Addition of RBC buffer and ethanol allowed RNA binding to MiniElute spin columns. After washing steps, pure RNAs were eluted with 20 μl of RNase-free water. Total RNA concentrations were determined with a Nanodrop spectrophotometer (Fisher Scientific).

### miRNA reverse transcription and quantification

miRNAs were specifically reverse transcribed with TaqMan Advanced miRNA cDNA Synthesis Kit (Applied Biosystems). Depending on RNA concentration, 10 ng or 2 μl total RNA were used as starting material for each poly(A) tailing reaction, followed by adaptor ligation and reverse transcription. We chose not to perform the last preamplification reaction in order to avoid eventual amplification bias.

The expression level of 752 miRNAs was screened by real-time PCR with TaqMan Advanced miRNA Human A and B Cards (Applied Biosystems A31805). cDNAs were diluted 1:10 with 0.1X TE buffer, then mixed with water and TaqMan Fast Advanced Master Mix 2X (Applied Biosystems) and 100 μL of this mix was loaded in each fill reservoir of two array cards. Real-time PCR reactions were run on a QuantStudio 7 Flex Real-Time PCR System (Applied Biosystems).

### mRNA reverse transcription and quantification

40 ng total RNAs were reverse transcribed with SuperScript IV Reverse Transcriptase (Invitrogen) and random hexamers in 30 μl total reaction volumes. cDNAs were diluted with water and 266 pg of cDNA was used in each 20 μl-PCR reaction in 96-well plates. Gene expression was quantified by real-time PCR with marmoset specific TaqMan Gene Expression Assays (Applied Biostems) and TaqMan Fast Advanced Master Mix (Applied Biosystems) on a QuantStudio 7 Flex Real-Time PCR System (Applied Biosystems).

### Data analysis

Data were revised and analyzed using ThermoFisher Scientific Digital Science online tools (thermofisher.com/fr/en/home/digital-science.html). Relative quantification was performed with the ΔCt method.

368 miRNAs were robustly amplified in more than 75% of the samples and were considered for subsequent analysis. From them, 103 were shared by the NeuN+ and NeuN-fractions. ΔCt values were obtained by global normalization method. qPCR results were first normalized (using global mean normalization method) and then transformed to relative expression levels via the 2^-ΔCt^ equation.

Four references genes were used as endogenous control genes (POLR2A, TBP, HPRT1, PGK1). ΔCt values were obtained by subtracting the mean Ct value of these 4 control genes to the Ct value of each target gene.

To obtain potential mRNA targets, we applied miRNet algorithm [28]. From the list of potential targets (546), we refined our list to 25 targets using the following criteria: i) mRNAs targeted by, at least, 1 of the 6 miRNAs dysregulated in area 32; and ii) the potential targets have been confirmed using an alternative prediction algorithm (Target Scan). Among them, 15 mRNAs contained target sequences for 3 miRNAs (high probability targets) and another 10 only 1 putative binding site (low probability targets). As a control, we also selected 10 reference genes bearing no binding sequence for any of the 6 miRNAs.

For behavioral experiments, EFA in HI test or snake model tests were extracted as previously described [29].

### Statistics

All values were represented with the mean ± SEM unless indicated. Statistical analysis was performed using XLStat (PCA), GraphPad 7.0 (ANOVA, correlation analysis and post-hoc tests) and R (PCA and regression analysis). A significance threshold of a 0,05 was used in all experiments.

No statistical methods were used to determine the sample sizes, but the number of experimental subjects is similar to sample sizes routinely used in our laboratory and in the field for similar experiments. We first tested the normal distribution of our data set using the Shapiro-Wilks and Kolmogorov-Smirnov tests. All data were normally distributed and variance was similar between groups, supporting the use of parametric statistics.

For all PCA analysis, we used Pearson correlation and n standardization. For the discrimination of NeuN+ and NeuN-fractions, we considered the expression levels of 103 shared miRNAs in both fractions. Although each fraction expresses 225-250 miRNAs, we displayed in all figures PCA analysis obtained from the common pool of 103 miRNAs. Apart from the first two, none of the remaining principal components (we analyzed 15) show any correlation to the cell fraction, region or genotype. The list of the 103 miRNAs is shown in Suppl Table 4.

For the analysis of miRNAs differences linked to SLC6A4 variants, we first performed a 2-way ANOVA on the relative expression levels of the 103 miRNAs shared by NeuN+ and NeuN-fractions in each cortical region. To limit type II error intrinsic to multiple comparison corrections, we reduced the dimensionality of our dataset to the most relevant miRNAs raised by our PCA analysis. Thus, we selected 15 miRNAs showing the strongest contribution to PC2 (the main dimension associated to the genetic polymorphisms in area 32) and 10 from the PC1. Although representing a small fraction of the analyzed miRNAs, these 25 miRNAs contribute to almost 40% and 30% of PC2 and PC1, respectively. We compared the expression levels of these 25 miRNAs in area 32 and 25 using 1-way ANOVA and Bonferroni’s post-hoc test to adjust for multiple comparisons (AC/C/G versus CT/T/C). The list of this miRNAs as well as the adjusted p values are presented in Table 1.

**Table 1.** Statistical analysis of expression levels of top 25 miRNAs from PCA (Oneway ANOVA adjusted for multiple comparison with Bonferroni’s correction). The first 10 miRNAs correspond to PC1 (gray shading) and the last 15 to PC2 (no shading).

| miRNA | Area 25 (AC vs CT)<br>Adjusted p value | Area 32 (AC vs CT)<br>Adjusted p value |
| --- | --- | --- |
| miR-628-3p | >0.999 | >0.999 |
| miR-645 | 0.4508 | >0.999 |
| miR-129-1-3p | 0.2080 | 0.6278 |
| miR-144-3p | >0.999 | >0.999 |
| miR-497-5p | 0.9585 | 0.6780 |
| miR-195-5p | >0.999 | 0.6839 |
| miR-196a-5p | 0.4076 | 0.5621 |
| miR-1260a | 0.1224 | 0.5981 |
| miR-125b-5p | 0.4031 | **0.0196** |
| miR-551a-5p | 0.4356 | >0.999 |
| miR-9-5p | 0.9891 | **0.0475** |
| miR-200a-3p | >0.999 | 0.1737 |
| let-7d-5p | >0.999 | **0.0208** |
| miR-26a-5p | >0.999 | 0.4908 |
| miR-190a-5p | >0.999 | **0.0032** |
| miR-29a-3p | 0.9459 | 0.0688 |
| let-7a-5p | >0.999 | 0.1181 |
| miR-133a-5p | >0.999 | 0.8187 |
| miR-124-3p | 0.3564 | 0.1234 |
| miR-378a-3p | >0.999 | 0.0554 |
| miR-376a-3p | 0.5647 | >0.999 |
| miR-320a | 0.9693 | >0.999 |
| miR-525-3p | 0.3089 | **0.0019** |
| miR-125a-5p | >0.999 | **0.0013** |
| miR-181c-5p | 0.8613 | >0.999 |

For the analysis of downstream targets, we analyzed the relative abundance of the mRNAs in area 25 and 32 using 2-way ANOVA followed by Tukey test.

Correlations were calculated using the Pearson correlation coefficient with 2-tailed analysis. Individual p values were adjusted for multiple comparison using the Holm-Sidak correction.

## RESULTS

### microRNA profiling in the marmoset cortex discriminates between NeuN^+^ and NeuN^-^ cells across cortical areas

In order to investigate whether previously described genetic variants in marmoset 5-HTTLPR result in miRNAs alterations that could be linked to behavioral responses, we first validated an experimental approach previously applied to human samples [30] (Fig. 1a). Brains from genotyped and behaviorally phenotyped marmosets were sliced and punches from selected brain regions were harvested. After nuclear isolation, samples were FACS sorted into NeuN^+^ and NeuN^-^ cells (Suppl. Fig. 1a) and RNA extracted from each fraction. As expected, NeuN^+^ cells are enriched in neuron-specific markers (Grm7, Gabra1, Camk2) and deprived almost entirely of glial-associated genes (astrocyte, oligodendrocytes and microglia markers, Fig. 1b and Suppl. Fig. 1b). In contrast, NeuN^-^ cells express strong levels of astrocytes (Gfap, Aldh1l1 or Slc1a3), oligodendrocytes (Klk6, Plp1, Cnskr3) and microglial genes (Aif1) (Fig. 1B and Suppl. Fig. 1b). We also observe a low expression of neuronal genes in this fraction as it is known that a subset of neurons of the primate cortex are NeuN^-^ [30]. Similar profiles were obtained in samples from different cortical regions (area 17, 25 and 32) indicating that FACS sorting is a reliable method to enrich neurons from different marmoset cortical areas for transcriptomic analysis.

**Figure 1.**
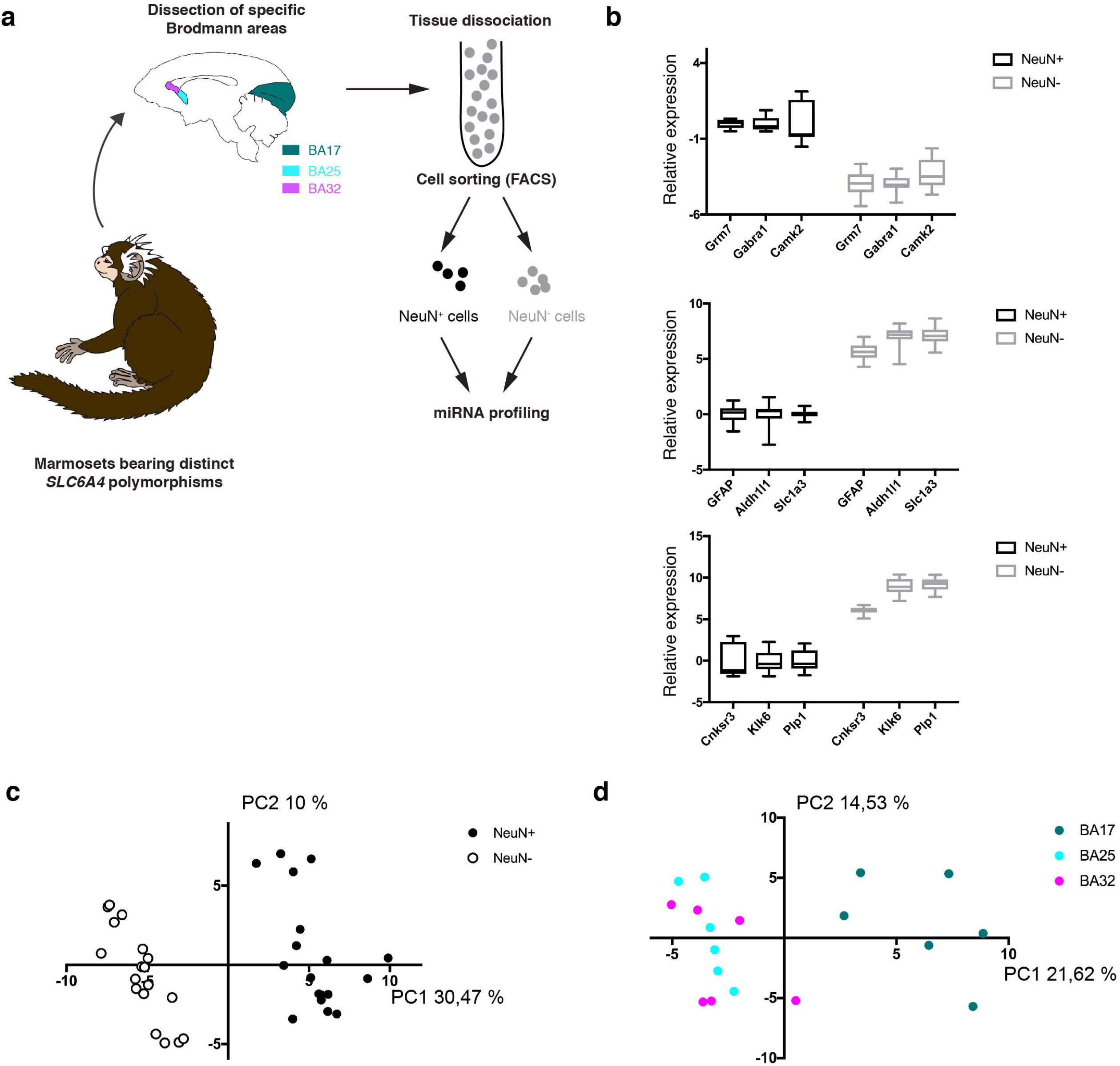
Schematic representation of experimental protocol and validation steps. a) Experimental protocol includes the genotypic and phenotypic characterization of the marmosets. After sacrifice, brains were frozen and sliced without fixation. RNA was extracted from punches of different cortical regions. Samples were previously submitted to nuclear isolation, NeuN staining and FACS sorting. b) Expression of neuronal (top panel), astrocytic (middle panel) and oligodendrocytic (bottom panels) markers in NeuN+ and NeuN-fractions confirms the efficiency of the FACS sorting strategy. c) miRNA profiling enables differentiation of NeuN+ and NeuN-subsets. Using PCA on 103 shared miRNAs, NeuN+ and NeuN-nuclei clearly segragate across the PC1 axis. d) Discrimination of regional differences based on miRNAs levels. PCA analysis clearly distinguishes the visual cortex (positive values) from the highly associative areas of the vmPFC (negative values).

Although miRNA expression in different brain cell types remains largely unexplored, we hypothesize that, given the differences in cell composition, miRNAs signatures present in NeuN^+^ and NeuN^-^ populations should be dramatically distinct. Using miRNA quantitative PCR, we profiled 754 miRNAs in our samples and found that more than 100 miRNAs were shared by both subpopulations. Focusing on this common pool of miRNAs, we performed a principal component analysis (PCA) in NeuN^+^ and NeuN^-^ nuclei coming from 6 different marmosets. As shown in Fig. 1c, NeuN^+^ and NeuN^-^ samples formed separated clusters across the major PC1 axis confirming that, even considering only those miRNAs whose expression is shared, miRNAs profiles readily distinguish both fractions.

Recent work revealed important regional differences in gene expression across the marmoset cortex [31]. We next investigated whether, similarly, miRNA profiling might be sensitive enough and detect such regional variations. For that purpose, we examined 3 cortical areas; on one hand, we profiled the primary visual cortex (corresponding to Brodmann area 17) as an example of sensory region endowed with specific cytoarchitectonic and functional features (e.g. expanded layer IV, strong myelination, major inputs from the thalamus). On the other hand, we considered two high-order association areas within the vmPFC (Brodmann area 25 and 32) whose activity has been shown to be consistently dysregulated in affective disorders [32]. A clear segregation between sensory and association areas in terms of miRNAs signature was reconstructed using principal component analysis (PCA) in NeuN^+^ cells (Fig. 1d). Whilst samples from the visual cortex clustered together on one side of the PC1 axis, the two vmPFC regions appeared intermingled on the other side of the PC1. Accordingly, a number of miRNAs (Suppl. Fig. 1d) are differentially expressed in the visual cortex. Such regional pattern cannot be observed in the NeuN^-^ fraction (Suppl. Fig. 1c) suggesting that anatomo-molecular differences largely arise from neurons. Together, these findings support the notion that miRNAs profiling is a powerful method to uncover molecular differences in the brain.

### microRNA profiling uncovers region-specific molecular differences in marmosets bearing different SLC6A4 variants/haplotypes

Since half of the 6 marmosets analyzed here beared each of the two most frequent *SLC6A4* haplotypes (AC/C/G versus CT/T/C), we next sought to determine whether miRNA profiling could unveil molecular differences related to those polymorphisms ^34^. Fig. 2a depicts the PCA analysis in the NeuN^+^ fraction according to the genetic variant of each animal as well as the anatomical region. While there is no obvious link in area 17 or 25, each haplotype segregated into two independent clusters across the PC2 axis in area 32. These results suggest that this region might be specifically affected by *SLC6A4* polymorphisms. In order to confirm this genetic effect, we carried out a 2-way ANOVA on the miRNA expression levels in each region. We found that only area 32 exhibited a significant effect of genotype on the miRNA expression (F(1, 4)=20.27; p=0.0108) whereas this was not observed in areas 17 (F(1, 4)= 1.973; p=0.2328) and 25 (F(1, 4)=3.244; p=0.146).

**Figure 2.**
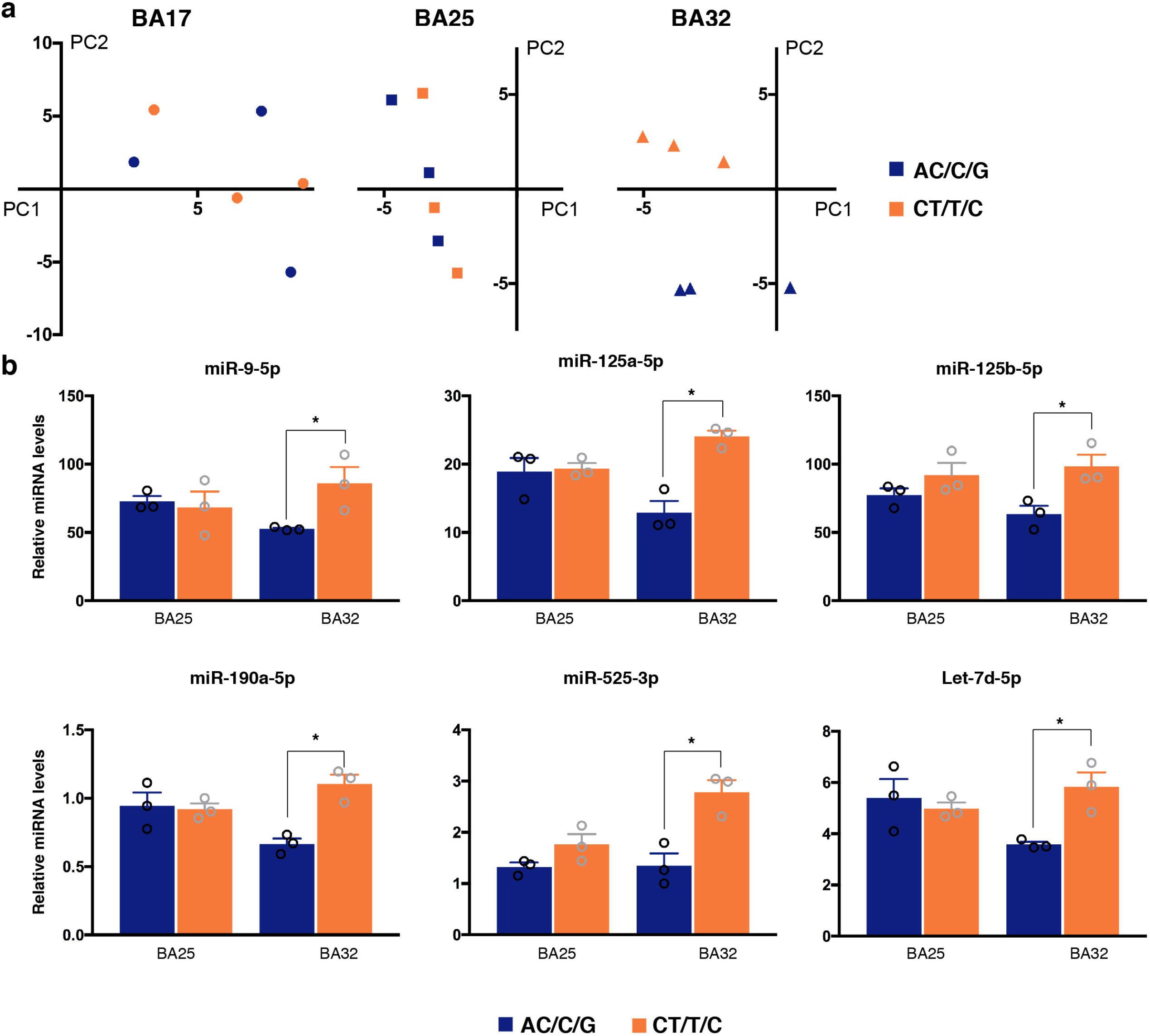
SLC6A4 polymorphisms (AC/C/G and CT/T/C) alter miRNA signature in area 32. a) PCA analysis on miRNAs levels in NeuN+ nuclei shows genotype-linked differences in area 32. b) miRNAs differentially expressed in area 32 in AC/C/G and CT/T/C marmosets (One way ANOVA followed by Bonferroni’s correction for multiple comparisons, * p<0.05).

To determine which miRNAs are significantly and specifically deregulated in area 32 (differentially expressed miRNAs, DEmiRs), we focused on the 25 miRNAs that showing the strongest contribution to the first two components in the PCA (Table 1, see also materials). We found that 6 out these 25 miRNAs (miR-9-5p miR-125a-5p, miR-125b-5p, miR-190a-5p, miR-525-3p and let-7d-5p,) exhibited differential expression in area 32 but not in the closely related area 25 (Fig. 2b).

As stated above, we found a number of miRNAs (miR-195, miR-221, miR-222 and miR-497) to be differentially expressed in the visual cortex compared to vmPFC (Suppl. Fig. 1 and 2). To further confirm the specificity of area 32 miRNA changes, we also examined whether SLC6A4 polymorphisms impinged on the expression of miRNAs associated with visual cortex. As shown in Suppl. Fig 3, none of these miRNAs showed any difference across the genotypes either in the visual cortex or in the vmPFC, suggesting that there exist miRNAs whose expression is altered in precise regions and in a genotype-dependent manner. Overall, our observations confirm that miRNAs could reliably uncover molecular differences in the marmoset cortex and indicate *SLC6A4* polymorphisms selectively alter miRNA signatures in area 32.

### Genotype-specific changes of DCC expression in area 32

miRNAs regulate gene expression post-transcriptionally. We reasoned that miRNAs alterations in area 32 would result in significant changes in downstream target transcripts. To identify those targets and thus further validate our miRNAs signatures, we carried out a network analysis of the area 32 deregulated miRNAs using miRNet [28] (Fig 3a). Using additional criteria (see Material and Methods), we selected 25 target genes (10 low probability targets and 15 high probability targets) as well as 10 control transcripts for further expression analysis in area 25 and 32. We found no difference across haplotypes in the expression of any low probability target or reference targets in area 32 (Suppl. Fig 3). Among the 15 high probability targets (Suppl. Fig 3), only DCC was found to be differentially expressed depending on *SLC6A4* polymorphisms (Fig 3). Although DCC transcript contains putative sequences for miR-9-5p, let7-5p and miR-190, the observed downregulation was moderate, in agreement with the contention that miRNAs fine-tune gene expression [33, 34]. As expected, levels of DCC and those of miR-9-5p, let7-5p and miR-190 miRNAs were inversely correlated. Thus, CT/T/C marmosets show reduced DCC and high levels of those miRNAs compared to AC/C/G animals (Fig. 2b). Finally, we observed no significant alteration of any transcript including DCC in area 25 (Fig. 3 and Suppl. Fig 3) arguing again for the anatomical specificity of molecular alterations associated with *SLC6A4* polymorphisms.

**Figure 3.**
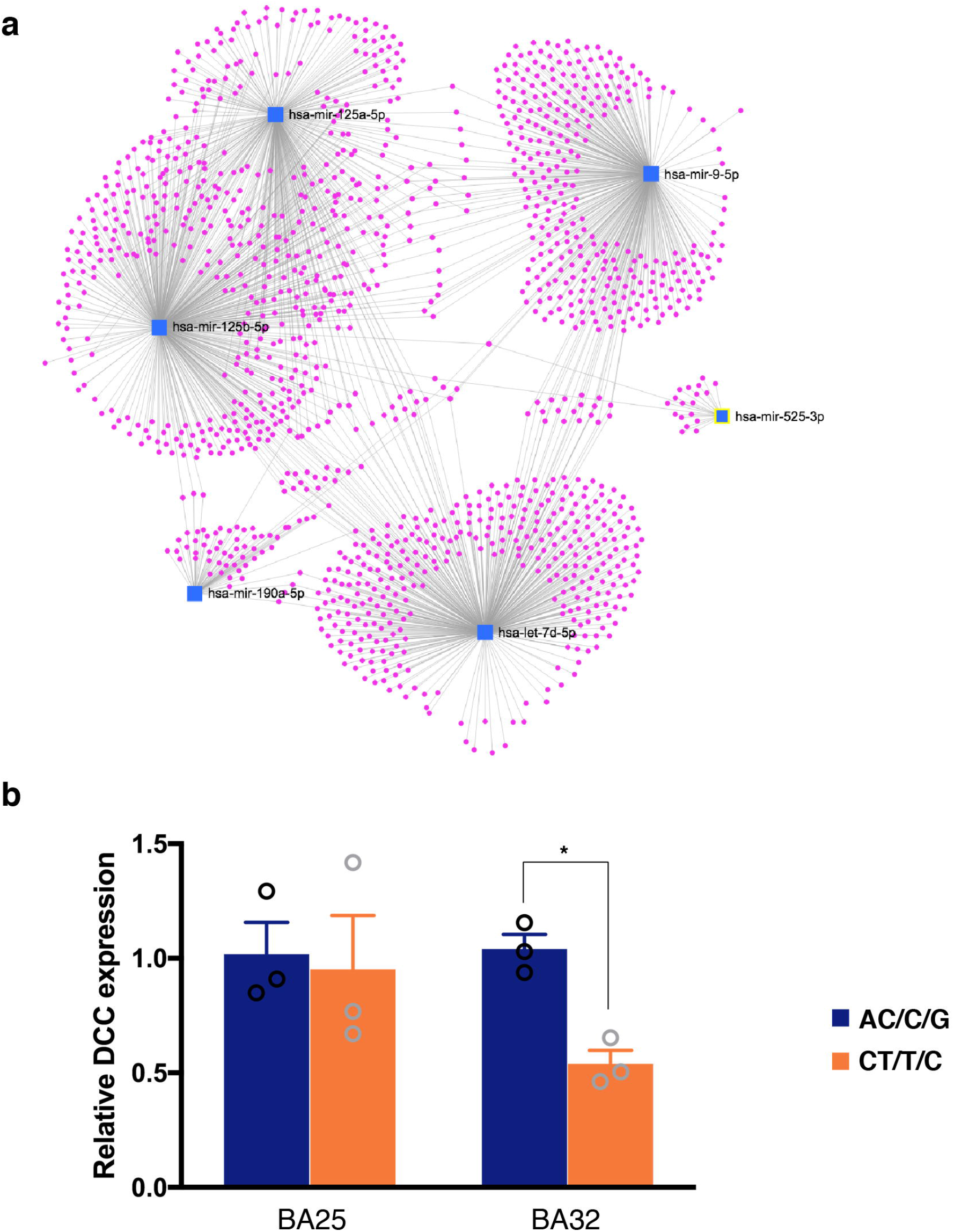
Target mRNAs dysregulated in area 32. a) Network analysis using miRNAs deregulated in area 32. b) Changes in DCC expression in area 32 are related to SLC6A4 variant. DCC was found to be significantly decreased in area 32 of marmosets bearing CT/T/C haplotype (2 way ANOVA followed by Tukey’s correctionfor multiple comparisons, * p<0.05,)

### Molecular alterations in area 32 correlate with behavioral response to uncertain threat

It has been previously shown that *SLC6A4* polymorphisms strongly influence anxiety-like behavior in response to uncertainty in the human intruder test (HI-test) but do not alter evoked fear-like behavior in the more certain context of the snake test (ST) [17, 26]. Using exploratory factor analysis (EFA), a recent study demonstrated that a single factor in the EFA explained behavior on the HI test whereas two factors were necessary to describe behaviors elicited on the ST [29].

We reasoned that, if relevant, the molecular alterations identified in area 32 might correlate with behavioral responses in the anxiety-related HI-test but not the fear-related snake test. We therefore performed a correlation analysis on the levels of miRNAs deregulated with the EFA score for HI-test. We observed a clear negative correlation for miR-125a-5p, miR-125b-5p, miR-525-3p and let-7d-5p in area 32 (Fig. 4), showing a *R^2^* coefficient ranging from 0.68 to 0.94. A significant association was observed for two of them (miR-525-3p and let-7d-5p) after correcting for multiple comparisons. Remarkably, DCC contents also showed a significant but inverted correlation to the behavioral score (*R^2^*=0.8994). In sharp contrast, levels of the same miRNAs and DCC in area 25 exhibited no correlation with the HI test EFA (Fig. 4) arguing for the specificity of our findings. Moreover, behavioral specificity was indicated by the finding that the two behavioral scores in the ST were not correlated with any of these miRNAs in area 32 (Suppl. Fig. 4), altogether supporting the notion that the molecular alterations we found in area 32 are selectively related to the differential behavioral response to uncertain threat in marmosets bearing the different *SLC6A4* variants.

**Figure 4.**
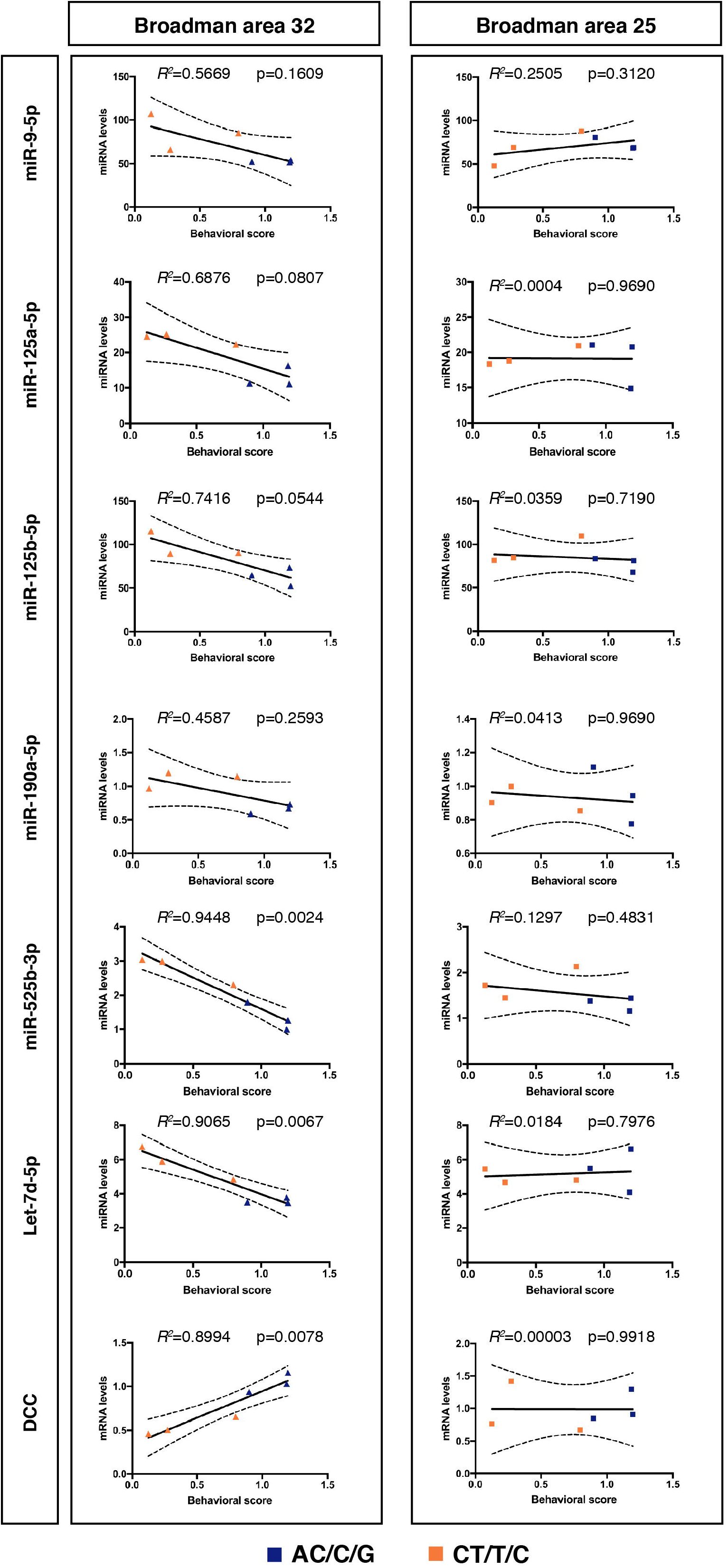
Correlation between miRNA and DCC levels in area 32 (left panels) or 25 (right panels) and behavioral response in the human intruder test. Individual p values are adjusted for multiple comparison using the Holm-Sidak correction.

Finally, recent work has suggested that the effectiveness of different drugs to dampen anxiety-behavior towards the intruder differs in marmosets homozygous for AC/C/G versus CT/TC variants [17, 19]. We asked whether these differential responses to citalopram (10 mg/kg) or the selective 5-HT2A antagonist, M100907 (0.3 mg/kg) were significantly correlated with miRNAs levels. We focused on the average distance the animals moved in response to the human intruder as this parameter is influenced by both pharmacological treatments [17]. As shown in Fig. 5, whilst only miR-9-5p levels in area 32 were specifically associated with the citalopram response (*R^2^*=0.7809), M100907 effects were correlated instead with those of miR-525-3p (*R^2^*=0.7562). No miRNAs in area 25 showed any correlation with the responses to citalopram or M100907 treatment supporting the anatomical segregation of molecular changes (Suppl. Figure 5). Together, these observations reveal the intimate links between area 32 molecular composition (specific miRNAs and DCC) and its function in the elaboration of complex behavioral responses towards uncertain threats.

**Figure 5.**
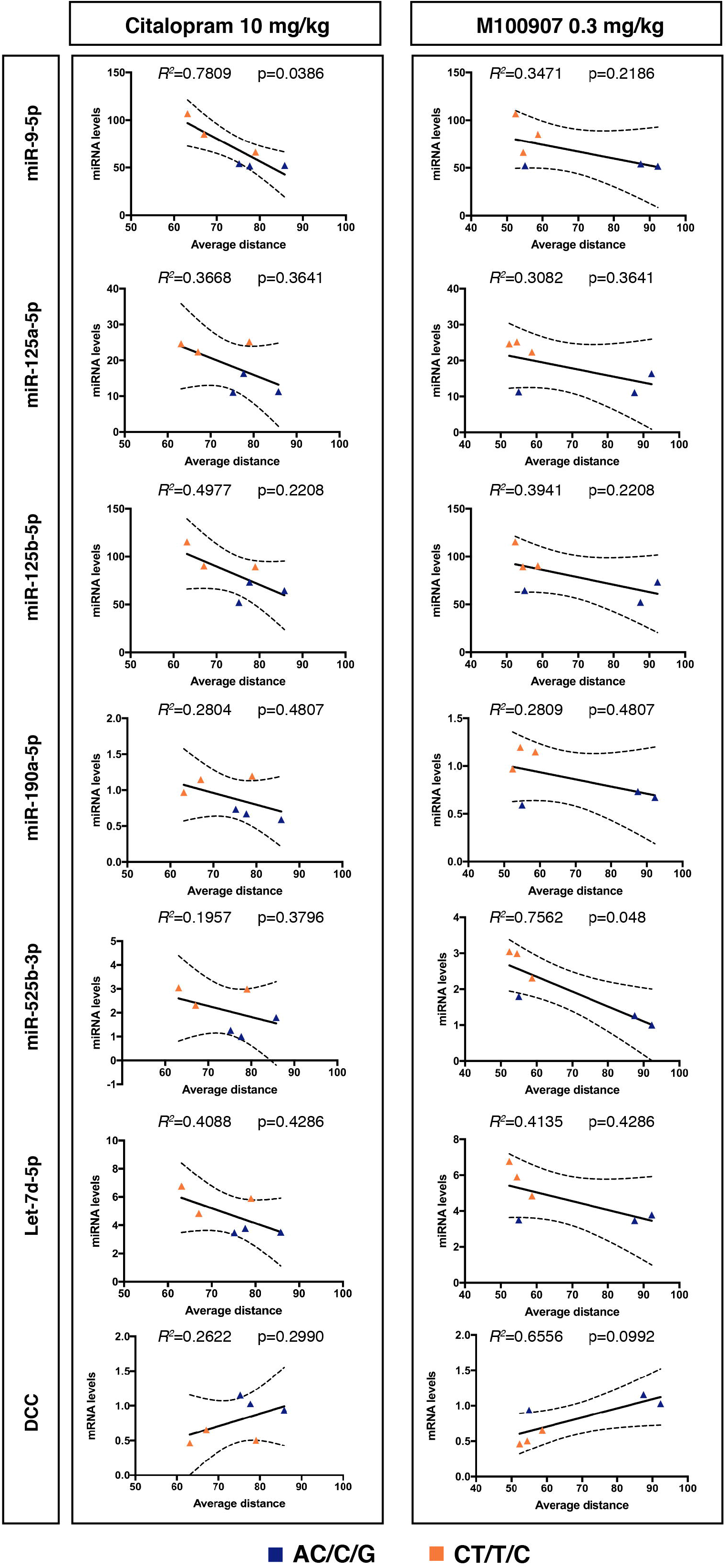
Correlation between miRNA and DCC levels in area 32 and average distance during the intruder phase after an acute injection of citalopram (left panels) or the 5-HT2a antagonist M100907 (right panels). Individual p values are adjusted for multiple comparison using the Holm-Sidak correction.

## DISCUSSION

Here, we setup a FACS-based approach never applied before in non-human primates to assess miRNA expression in different cell subsets of behaviorally phenotyped animals. Our results show that: i) miRNAs are dramatically different across cell types and cortical regions; ii) *SLC6A4* polymorphisms impinge selectively on miRNA signatures in area 32 of vmPFC; iii) levels of specific miRNAs as well as of the target gene, DCC, correlate with anxiety-like behavior in response to uncertain threat in the intruder test; and iv) the responsiveness to anxiolytics/antidepressants is also associated with specific miRNAs. Although based on a limited number of samples, our results highlight the refinement of genetic-molecular-behavioral correlations in a primate.

Research in biological and molecular psychiatry is confronted by the enormous differences between the human brain and those of experimental animals most commonly used (mice, fish or fly). Non-human primates, especially marmosets, might represent a more appropriate model. First, in terms of genome, primates are evolutionary closer to humans and share most of the coding and non-coding sequences. Thus, 5-HTTLPR genomic arrangement is highly conserved in primates but does not exist in other vertebrates such as mouse or zebrafish. Similar to that reported in humans [4, 35], naturally occurring polymorphisms in the marmoset *SLC6A4* promoter region are linked to anxious-like behaviors [17, 19] highlighting the conservation of genomic structure and function. Second, brain anatomy is more similar to humans than that found in these other species, especially in those areas related to affective disorders [36]. Finally, in molecular terms, it has been shown that the repertoire of non-coding RNAs (ncRNAs) has expanded across evolution, and multiple clinical studies support the idea that primate-restricted ncRNAs contribute to psychiatric conditions including depression/anxiety [23, 37]. Our findings in non-human primates provide additional evidence to support this notion. Thus, miR-525b-3p, one of the miRNAs whose levels in vmPFC area 32 depend on the SLC6A4 genetic variants, is an evolutionary recent miRNA as shown by the exact sequence conservation between human, gorilla, and chimpanzee (Suppl. Fig 6a). A less conserved sequence is found in other old-world (orangutan, baboon and macaques) and new-world monkeys (marmosets and squirrel monkeys) but not in prosimians (mouse lemur). Similarly, miR-190-5p binding sequences in DCC transcript are conserved across primates but mutated in rodents such as rats or guinea pig (Suppl. Fig 6b)

In light of our results, an important mechanistic question is how genetic variants in SLC6A4 affect miRNAs. Considerable evidence suggests that miRNAs have a pivotal role in conferring robustness to biological processes [38] and thus modulate gene x environment interactions [39]. In the brain, this refers to the ability to maintain a function in spite of genetic or environmental fluctuations. An appealing hypothesis is that, in response to genetic variation (i.e. 5-HTTLPR S allele in humans or AC/C/G in marmosets), miRNAs might provide a molecular mechanism to limit the functional impact of enhanced sensitivity to negative environmental events. Our results suggest that this might only occur in specific networks particularly vulnerable to negative environmental influences such as those involving vmPFC area 32. Area 32 is a key brain region involved in decision making in conflict environments as shown by studies of approach-avoidance decision making in primates [40] and, along with dorsolateral prefrontal cortex, with which it shares bi-directional connections, is dysregulated in depression [41]. Interestingly, a recent human study using PET scan suggested marked regional differences in the expression of key elements of serotonin neurotransmission within the vmPFC [8]. According to this work, area 32 showed considerably lower levels of the serotonin transporter than area 25 providing a rationale for the differential impact of the genetic polymorphisms on those two regions.

Our work has potential important implications for the treatment of affective disorders. First, miRNAs are short molecules endowed with an enormous therapeutic potential [42]. Here, we have identified a set of miRNAs whose levels are correlated to different behavioral outcomes in response to uncertainty measured by the HI test, a paradigm ethologically relevant for the investigation of stress-related disorders [43]. Second, as previously shown, miRNA-dependent fine-tuning of specific targets has been consistently involved in the pathophysiology of stress-related disorders [22, 44]. Our findings pinpoint the therapeutic potential of DCC, a well-established risk factor for depression [22, 34, 45]. Previous work demonstrated that DCC expression is under the control of miR-218 [33, 34]. Although we could not observe any difference in miR-218 associated to SLC6A4 variants (Suppl. Fig 6), our data support the implication of alternative miRNAs (namely, miR-9, miR-190a and let-7d) in the regulation of DCC. Similarly, Reynolds et al suggest that DCC operates on mesolimbic dopaminergic neurons [46]. Our observations open the possibility that DCC signaling is also active on serotoninergic terminals. Finally, since all molecular alterations described here are restricted to a discrete cortical region, our findings also raise a technical challenge of finding the most optimal delivery route for such therapeutic interventions. In summary, our work suggests that modulating certain molecular pathways (miRNAs or DCC) in a region-specific manner might unfold novel opportunities for precision medicine.

Accumulating evidence also indicates that miRNAs could be useful biomarkers in psychiatry as multiple studies have demonstrated that blood levels faithfully translate contents of miRNAs in the brain [22, 47]. In this regard, miRNAs have been shown to be particularly useful to evaluate the responsiveness to antidepressants [48]. Our observations expand this concept and suggest that specific miRNAs could be related to distinct pharmaco-therapy pathways. Thus, miR-9 levels predict the effects of serotonin re-uptake inhibitors on threat responsivity while miR-525 levels correlate with the response to 5-HT2a antagonists (Fig. 5). The influence of genetic variants on the levels of these miRNAs in a region-specific manner revealed here, highlight the need to integrate genetic, molecular, imaging and clinical data for personalized therapy.

## CONTRIBUTORS

ACR, AMS and EG conceived and designed the project. AMS performed the behavioral testing and sample preparation. NP performed miRNA and gene expression experiments. NP, AMS, DB and EG analyzed the data. ACR provided contribution to the interpretation of data. AMS, ACR and EG wrote the manuscript. NP, ACR, AMS and EG discussed the results, reviewed and edited the final manuscript.

## ACKNOWLEDGEMENTS

We would like to thank Stephane Robert from the AMUTICYT facility for his precious support for the FACS sorting. We also thank Catherine Lepolard for technical help.

