## Supplemental material for "Region-specific microRNA alterations in marmosets carrying SLC6A4 polymorphisms are associated with anxiety-like behavior"

**Supplementary Table 1. Subjects used in this study**

| Subject | Genotype | HI-test age,y | ST age, y | Age at death, y |
|---|---|---|---|---|
| Jetsam | CT/T/C | 1.71 | 2.01 | 5.66 |
| Sebastian | CT/T/C | 1.58 | 1.64 | 5.24 |
| Bob | CT/T/C | 2.44 | 2.53 | 6.13 |
| Bakerloo | AC/C/G | 2.43 | 2.51 | 6,11 |
| Merry | AC/C/G | 2.43 | 2.76 | 6.39 |
| Axel | AC/C/G | 2.86 | 3.21 | 6,83 |

**Supplementary Table 2. Behavioral parameters evaluated in the human intruder and snake test.**

| Human Intruder Test | Snake Test |
|---|---|
| Average height | Tsik-egg calls |
| Head and body bobs | Tsik calls |
| Time spent (back) | Stare count |
| Tse-egg calls | Stare duration |
| Locomotion | Locomotion |
| Time spent (front) | Average distance |
| Egg calls | Head-cock |
| Tsik calls | Egg calls |
| Tsik-egg calls | |

From Santangelo et al. 2016

**Supplementary Table 3. Primers used for sequencing**

| Primer name | Primer sequence |
|---|---|
| SLC6A4 repeat region forward | **CAGACAACCGTGTTCA TCTG** |
| SLC6A4 repeat region reverse | **GATTCTAGTGCCACCTAGAC** |
| Sequencing primer 1 | **AGCAGCACCTAACCCTCCTA** |
| Sequencing primer 2 | **TCCCCACTAGGCATTGCTAC** |

### Supplementary Table 4. List of miRNAs used for PCA analysis

let-7a-5p
let-7b-3p
let-7b-5p
let-7c-5p
let-7d-5p
let-7f-5p
let-7g-5p
let-7i-5p
miR-1-3p
miR-100-5p
miR-101-3p
miR-103a-3p
miR-107
miR-124-3p
miR-1248
miR-1255a
miR-125a-5p
miR-125b-5p
miR-1260a
miR-128-3p
miR-129-1-3p
miR-133b
miR-144-3p
miR-148a-3p
miR-153-3p
miR-15a-5p
miR-16-5p
miR-17-5p
miR-181a-5p
miR-181b-5p
miR-181c-5p
miR-185-5p
miR-190a-5p
miR-191-5p
miR-195-5p
miR-196a-5p
miR-197-3p
miR-199a-5p
miR-200a-3p
miR-200c-3p
miR-204-5p
miR-21-3p
miR-2110
miR-214-3p
miR-219a-5p
miR-22-3p
miR-24-3p
miR-26a-5p
miR-27b-3p
miR-29a-3p
miR-29b-3p
miR-29c-3p
miR-302b-3p
miR-302d-3p
miR-30b-5p
miR-30c-2-3p
miR-30d-5p
miR-320a
miR-323b-5p
miR-325
miR-338-3p
miR-339-3p
miR-342-3p
miR-361-3p
miR-361-5p
miR-374a-5p
miR-374b-5p
miR-376a-3p
miR-378a-3p
miR-380-3p
miR-423-5p
miR-448
miR-452-3p
miR-483-5p
miR-485-5p
miR-490-3p
miR-497-5p
miR-501-3p
miR-501-5p
miR-502-3p
miR-518f-3p
miR-524-3p
miR-525-3p
miR-548a-3p
miR-548e-3p
miR-548j-5p
miR-551a
miR-562
miR-576-3p
miR-593-3p
miR-606
miR-628-3p
miR-633
miR-645
miR-653-5p
miR-744-5p
miR-765
miR-9-3p
miR-9-5p
miR-92a-3p
miR-93-5p
miR-939-5p
miR-99b-5p

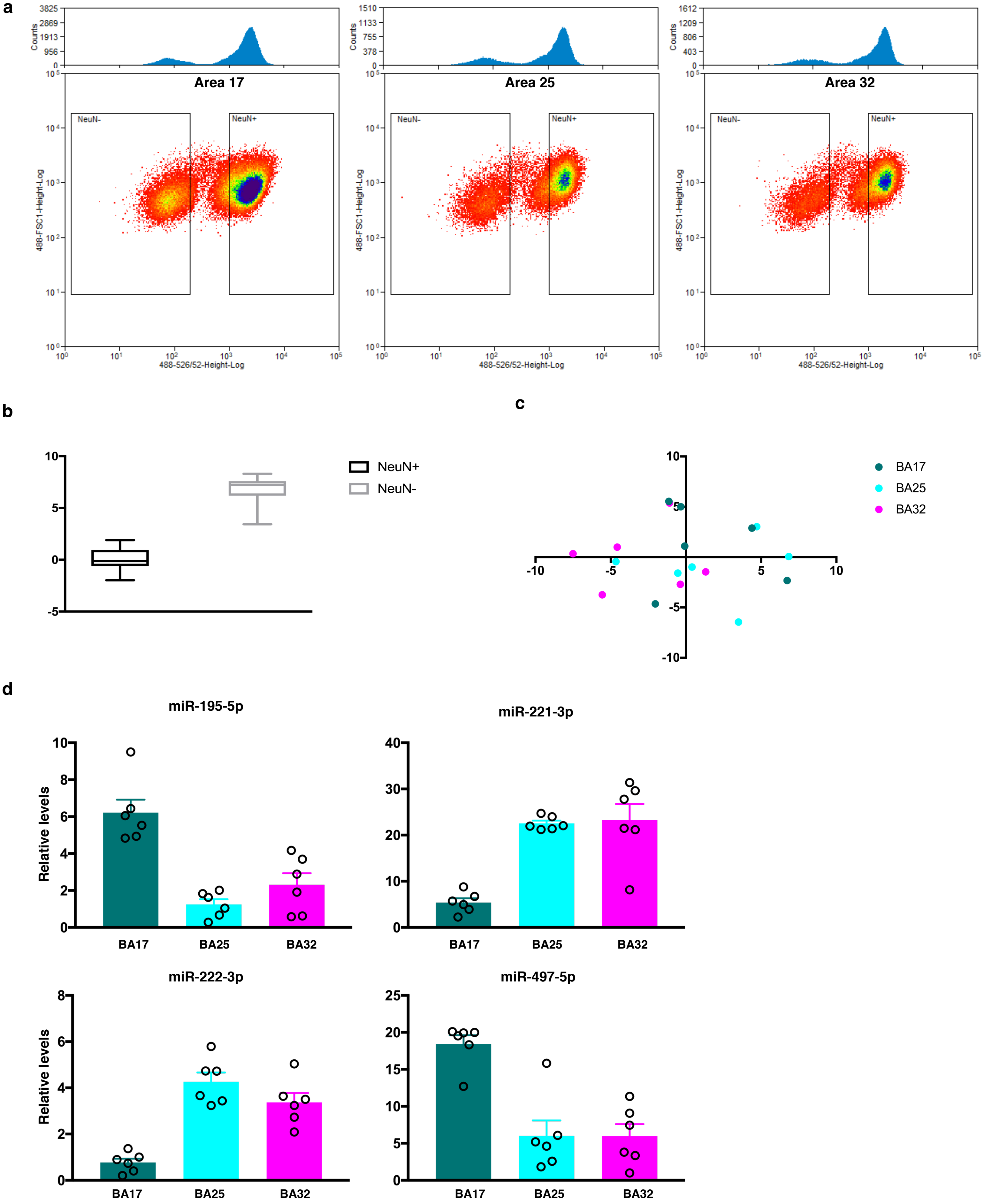


Supplementary Figure 1.
a) Representative example of the NeuN staining and windows used for FACS sorting.
b) Expression of microglial marker Aif2 in $NeuN^+$ and $NeuN^-$ fractions.
c) PCA analysis on miRNAs level in $NeuN^-$ nuclei shows no regional discrimination in this fraction.
d) Examples of miRNAs differentially expressed in the visual cortex. miR-221-3p/222-3p levels are significantly lower in BA17 compared to both areas of the vmPFC whereas miR-195-5p/497-5p are enriched in the visual cortex.

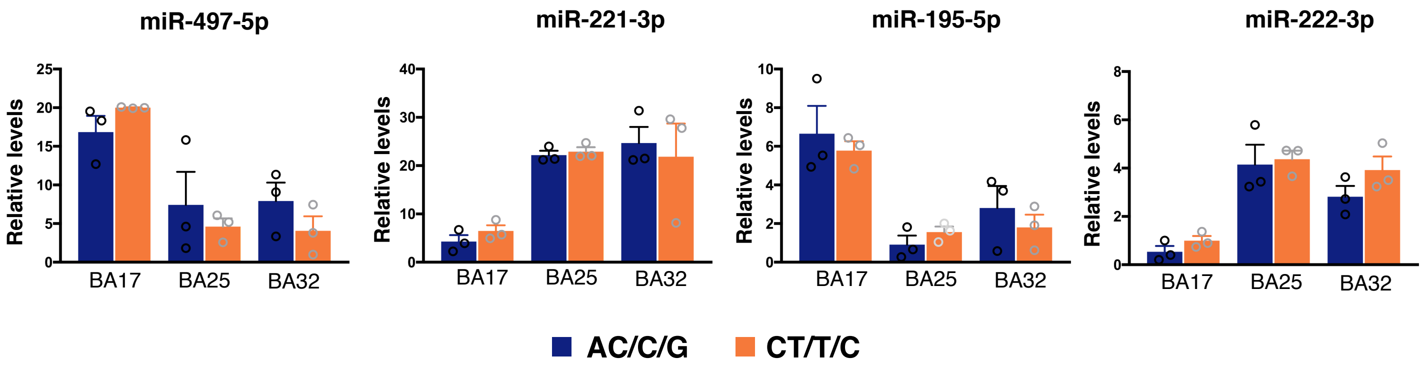


Supplementary Figure 2. miRNAs associated to the visual cortex show no differences in AC/C/G versus CT/T/C marmosets.

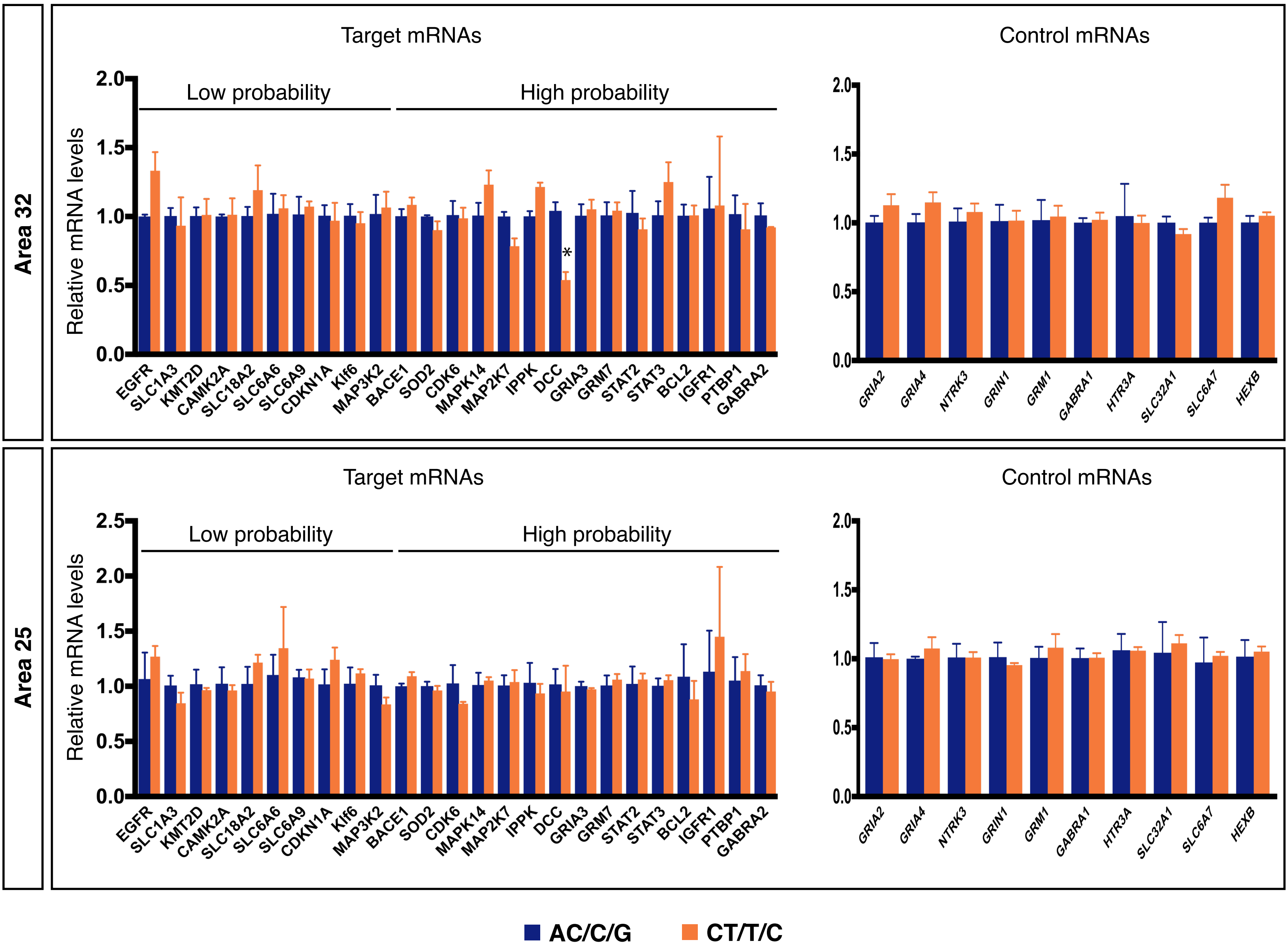


Supplementary Figure 3.
Expression of 25 target mRNAs identified by network analysis in area 32 (top) and 25 (bottom). Only DCC was altered in area 32. No significant differences were found in any of the tested transcripts in area 25. As a control, 10 reference genes were also quantified in both areas.

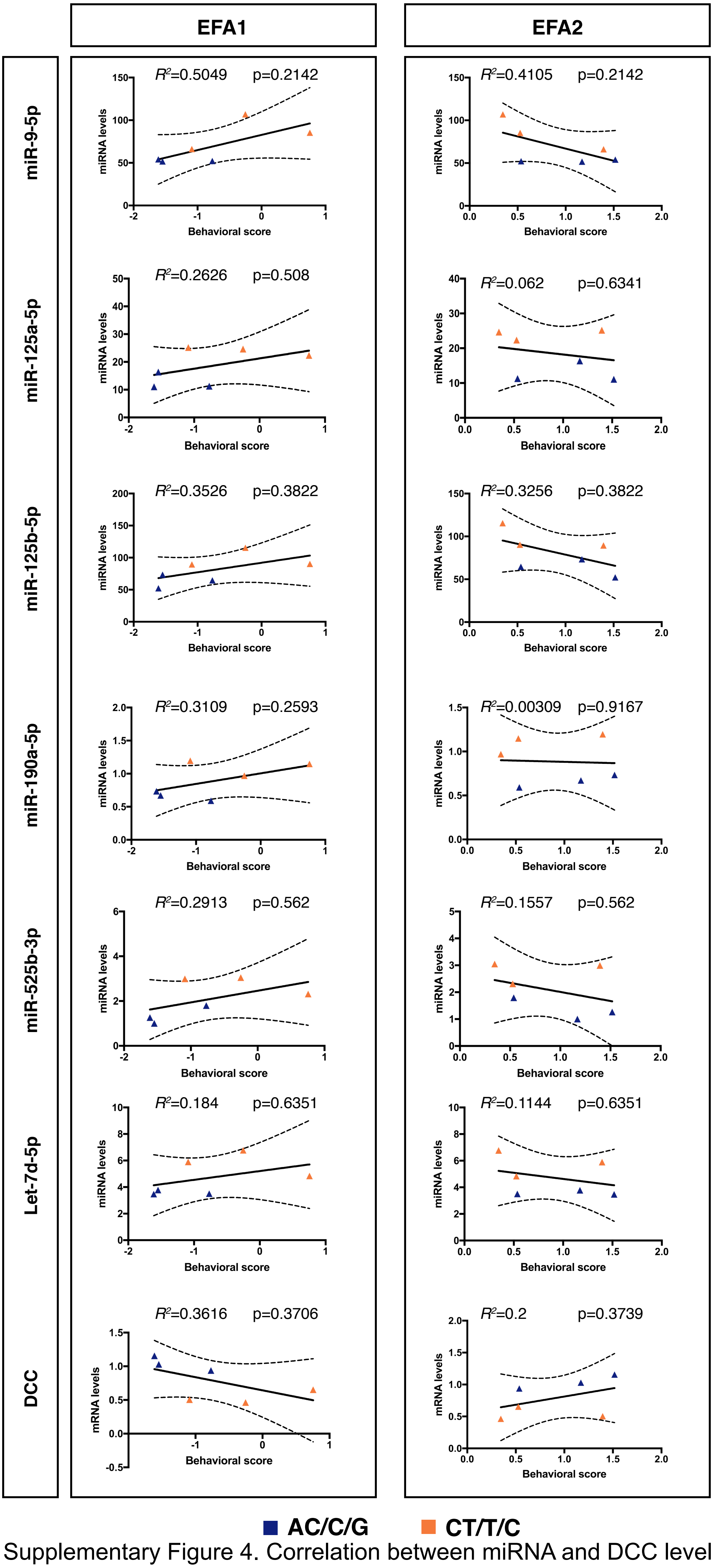


Supplementary Figure 4. Correlation between miRNA and DCC levels in area 32 and behavioral response in the snake test. Two EFA factors are considered in this analysis as previously reported (Quah et al. 2020).p values are adjusted for multiple comparison using the Holm-Sidak correction.

Popa et al Supplementary Figure 5

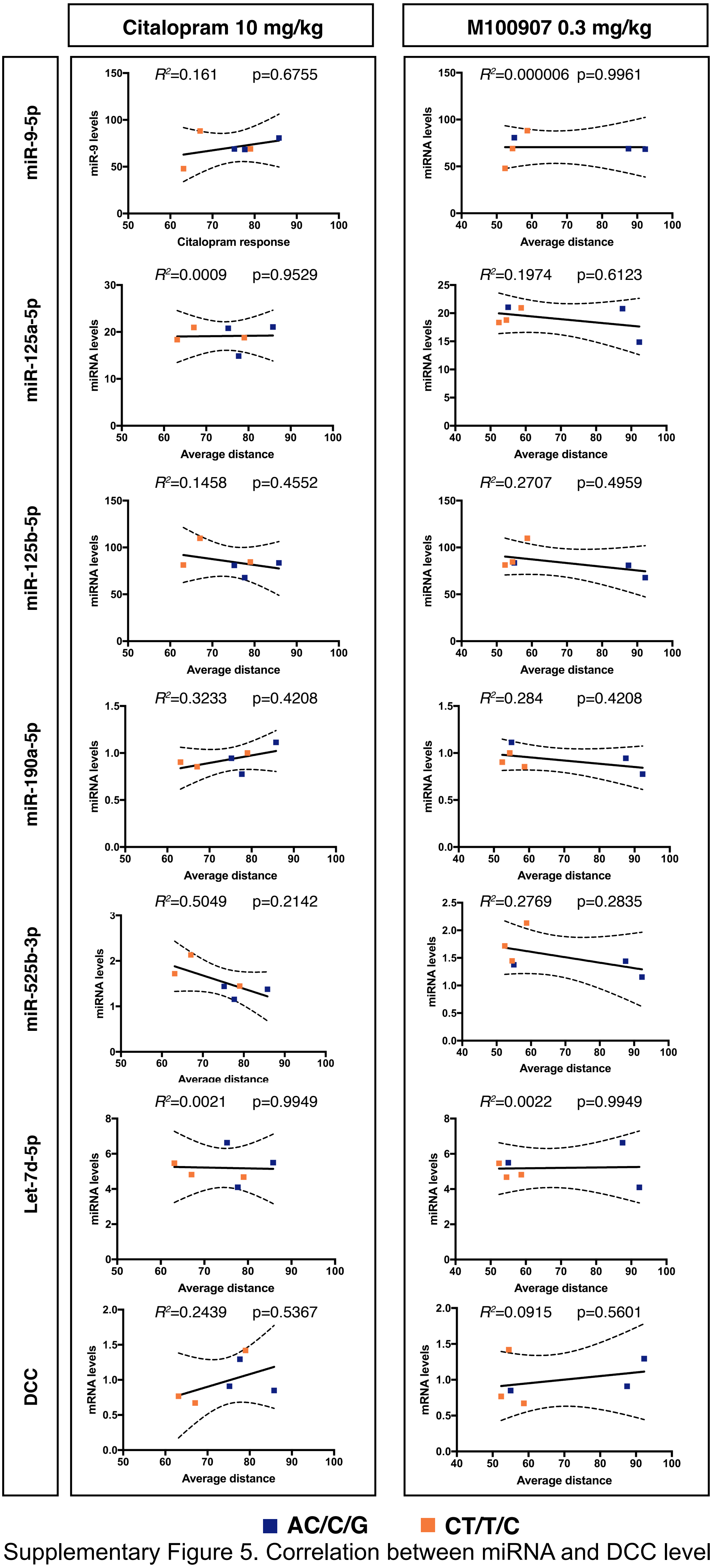


Supplementary Figure 5. Correlation between miRNA and DCC levels in area 25 and average distance during the intruder phase after an acute injection of citalopram (left panels) or the 5-HT2a antagonist M100907 (right panels). p values are adjusted for multiple comparison using the Holm-Sidak correction.

**a**

| | |
|---|---|
| Human | GAAGGCGCUUCCCUUUAGAGCG |
| Chimp | GAAGGCGCUUCCCUUUAGAGCG |
| Gorilla | GAAGGCGCUUCCCUUUAGAGCG |
| Orangutan | GAAGGCGCAUCCCUUUAGAGCG |
| Macaque | GAAGGCGCAUCCCUUUGGAGCG |
| Baboon | GAAGGCGCAUCCCUUUGGAGCG |
| Marmoset | GAAAGTGCUUCCCUUUAGAGTG |
| Squirrel monkey | GAACTTGCUUCCCUUUAGAGCG |

**b**

**miR-190a-5p binding site**

| | 4150 |
|---|---|
| Human | AUACCACACCCAUAUCAGCAAAUGAAUAUUACU |
| Marmoset | AUACCACAUCCAUAUCAGCAAAUGAAUAUUGCU |
| Rat | GUGCAGUGCCCAUAUGAGCAAAUGCGAGUAACU |

Supplementary Figure 6. a) Alignment of miR-525-3p in different primate species. b) Alignment of the 3' UTR DCC mRNA in humans, marmoset and rat to illustrate conservation/divergence of binding sites for miR-190a-5p.
